# The effect of population size on adaptation to fluctuating temperatures

**DOI:** 10.1101/2024.10.05.616761

**Authors:** Emmi Räsänen, Veera Nieminen, Pauliina A. M. Summanen, Mariana Villalba de la Peña, Peetu Makkonen, Kaisa Suisto, Tarmo Ketola, Ilkka Kronholm

## Abstract

Climate change exposes populations to more frequent periods of extreme temperatures and faster temperature fluctuations. Theoretical models suggest that different types of adaptations should occur in constant versus fluctuating environments of varying frequency. Furthermore, evolutionary adaptation to one environment may weaken the adaptations to alternative environments due to antagonistically pleiotropic alleles. However, fitness trade-offs are rarely observed in experiments and it has been hypothesized that the number and severity of trade-offs evolving in fluctuating environments might depend on population size. To evaluate whether specific types of adaptations evolve at fluctuating temperatures and how population size affects the evolution of trade-offs, we performed an evolution experiment with fission yeast (*Schizosaccharomyces pombe*). The small and large populations evolved for 500 generations at constant and fluctuating temperatures, after which the evolved strains competed against ancestral strains in respective selection environments and in alternative environments to detect trade-offs. We observed significant adaptation and maladaptation only to constant heat, but not to fluctuating temperatures. Overall, the population size did not have significant effects on adaptation capacity or trade-offs in alternative environments. Our results suggest that constant extreme temperatures may act as stronger selective pressures than temperature variation and that trade-offs are unlikely to constrain adaptation to fluctuating temperatures.

## 1 INTRODUCTION

Climate change is predicted to increase mean temperatures and also fluctuations in temperature (Meehl *et al*., 2000). For ectothermic organisms temperature affects many aspects of their fitness, and adapting to a new temperature regime can be challenging (Bennett and Lenski, 2007; Angilletta Jr., 2009). One of the most important factors determining the organisms’ capacity to adapt is a population size, as all else being equal, larger populations should have better mutational supply with more beneficial mutations of large effect size (Samani and Bell, 2010; Kristensen *et al*., 2020). This is alarming for the evolutionary potential of small and maladapted populations, as extreme temperature events and fluctuations can diminish population sizes and lower standing genetic variation, leading to greater genetic drift and extinction risks among species (Frankham, 2005; Willi *et al*., 2006). We need to better understand how populations of different sizes adapt to extreme temperatures and fast temperature fluctuations to be able to predict if populations can respond to climate change by evolutionary adaptation (Reed *et al*., 2011; Botero *et al*., 2015).

It has been suggested that the adaptation to fluctuating conditions may have a different genetic basis than adaptation to constant conditions (Ketola and Kristensen, 2017). Also, theoretical models predict that fluctuations of different frequencies should lead to specific outcomes of adaptation (Botero *et al*., 2015). If fluctuations are fast and happen within a generation, they should produce reversible plasticity or bet-hedging, and if fluctuations happen within tens of generations, developmental plasticity, possibly mediated by epigenetic mechanisms, can evolve (Botero *et al*., 2015; Kronholm, 2022). Furthermore, if environmental changes occur over hundreds of generations, then populations may track the environmental optimum by constantly adapting to it (Botero *et al*., 2015). Another alternative is that populations adapt by evolving a phenotype that tolerates fluctuations, or become generalists that do well across a wider range of temperatures (Kristensen *et al*., 2020).

Adaptation to fluctuating environments can also interact with population size. Recent studies have suggested that the adaptive cost and benefits in small and large populations should depend on the current variability in environmental conditions compared to the conditions during population’s evolutionary history (Chavhan *et al*., 2019, 2020, 2021). In general, populations that have evolved in fluctuating environments should have less fitness costs in alternative local environments, as under fluctuations, there is stronger selection against fitness costs due to antagonistically pleiotropic alleles (Bono *et al*., 2017). Conversely, when the conditions stay constant over a longer time, antagonistic pleiotropy can evolve more freely as selection is blind to the fitness costs in alternative environments (Bono *et al*., 2017). According to Chavhan *et al*. (2021), larger populations should have better access to mutations of large effect sizes, and these mutations may have stronger trade-offs in alternative environments. For example, when populations have evolved under constant conditions, larger populations should suffer from greater maladaptation when the environment changes (Chavhan *et al*., 2020). However, after evolution under fluctuating environment, larger populations should experience less trade-offs (Chavhan *et al*., 2021). As smaller populations are more likely to adapt by mutations of smaller effect sizes (Sniegowski and Gerrish, 2010; Chavhan *et al*., 2019), both the costs and benefits from the conditions during evolution should be minor and performance be more robust across alternative environments (Chavhan *et al*., 2021).

While there is some empirical evidence that the number and extent of trade-offs evolving in fluctuating environments might differ between large and small populations (Chavhan *et al*., 2021), no experiments have been done to investigate how population size interacts with trade-offs in thermal adaptation. We tested the hypotheses that different kinds of adaptations should evolve in constant and fluctuating environments depending on the speed of environmental change with respect to species’ generation time (Botero *et al*., 2015; Kristensen *et al*., 2020; Kronholm, 2022). We did an experimental evolution study of approx. 500 generations with fission yeast *Schizosaccharomyces pombe* populations that evolved either at constant temperatures or fluctuations of varying speed. Furthermore, we tested how population size affects thermal adaptation by including small and large populations for each temperature treatment. In particular, we wanted to study the fitness trade-offs in thermal adaptation and investigated the following questions: (1) Do populations have fitness costs at alternative temperatures if they have evolved at constant or fluctuating temperatures? (2) Do larger populations adapt more efficiently than smaller populations to constant or fluctuating temperatures? (3) Does population size affect the occurrence of trade-offs differently at constant and fluctuating temperatures? Our study is the first one attempting to directly test the effects of population size on adaptation to constant and fluctuating temperatures.

## 2 MATERIALS AND METHODS

### 2.1 Schizosaccharomyces pombe strains

In our evolution experiment, we used four *S. pombe* strains as ancestors. These represented all combinations of the two mating types *h*^+^ and *h*^−^, and two alleles of the *ade6* marker, *ade6^M^*^210^ and *ade6^M^*^216^. The *ade6* mutants are adenine auxotrophs, and when grown on plates with low adenine, their colonies develop different colours, as *ade-6^M^*^216^ colonies turn pink and *ade6^M^*^210^ mutants darker red (Forsburg and Rhind, 2006). We used *ade6* marker to distinguish different strains in competition experiments.

Since wild type *S. pombe* strains are capable of switching their mating type by somatic recombination (Hanson and Wolfe, 2017), we used strains for which the inactive *mat2* and *mat3* genes had been deleted, making them to have stable mating types. The full genotypes of the four ancestors were: *h*^+^ *(H1::hphMX)* Δ*mat2,3::LEU2 ade6^M^*^216^ *ura4^D^*^18^ *leu1*^32^, *h*^+^ *(H1::hphMX)* Δ*mat2,3::LEU2 ade6^M^*^210^ *ura4^D^*^18^ *leu1*^32^, *h*^−^ *(H1::hphMX)* Δ*mat2,3::LEU2 ade6^M^*^216^ *ura4^D^*^18^ *leu1*^32^, and *h*^−^ *(H1:: hphMX)* Δ*mat2,3::LEU2 ade6^M^*^210^ *ura4^D^*^18^ *leu1*^32^. Otherwise these strains are nearly isogenic and were tested to have similar growth rates. The strains were kindly provided to us by Dr. Bart Nieuwenhuis.

### 2.2 Evolution experiment

In the evolution experiment, populations of *S. pombe* evolved in five different temperature treatments; constant mean (34 ^◦^C, the optimal temperature), constant extreme (38 ^◦^C), fast, intermediate, and slow fluctuations (30–38– 30 ^◦^C). We chose the temperatures based on the thermal performance curve we did in a preliminary experiment for the ancestor strain *h*^+^ *ade6^M^*^216^ (Figure 1). The rates of fluctuations were chosen based on the models of Botero *et al*. (2015) and Kronholm (2022), and they are predicted to lead into different types of adaptations. At fast fluctuations one temperature cycle was completed in 24 h, at intermediate fluctuations in 13 days, and at slow fluctuations in 40 days. Based on preliminary experiments, the number of generations during one temperature cycle should be approx. 2.5, 32, and 100, for the fast, intermediate, and slow fluctuations respectively. In addition we had small (*N_e_* = 10^6^) and large (*N_e_* = 10^7^) populations in each temperature treatment. The population sizes were selected based on a literature review in which larger populations did not show fitness costs in fluctuating environments (Chavhan *et al*., 2021). The evolution experiment contained four replicate populations of the four ancestors, which with five temperature treatments and two population sizes gave in total 160 populations.

**Figure 1:**
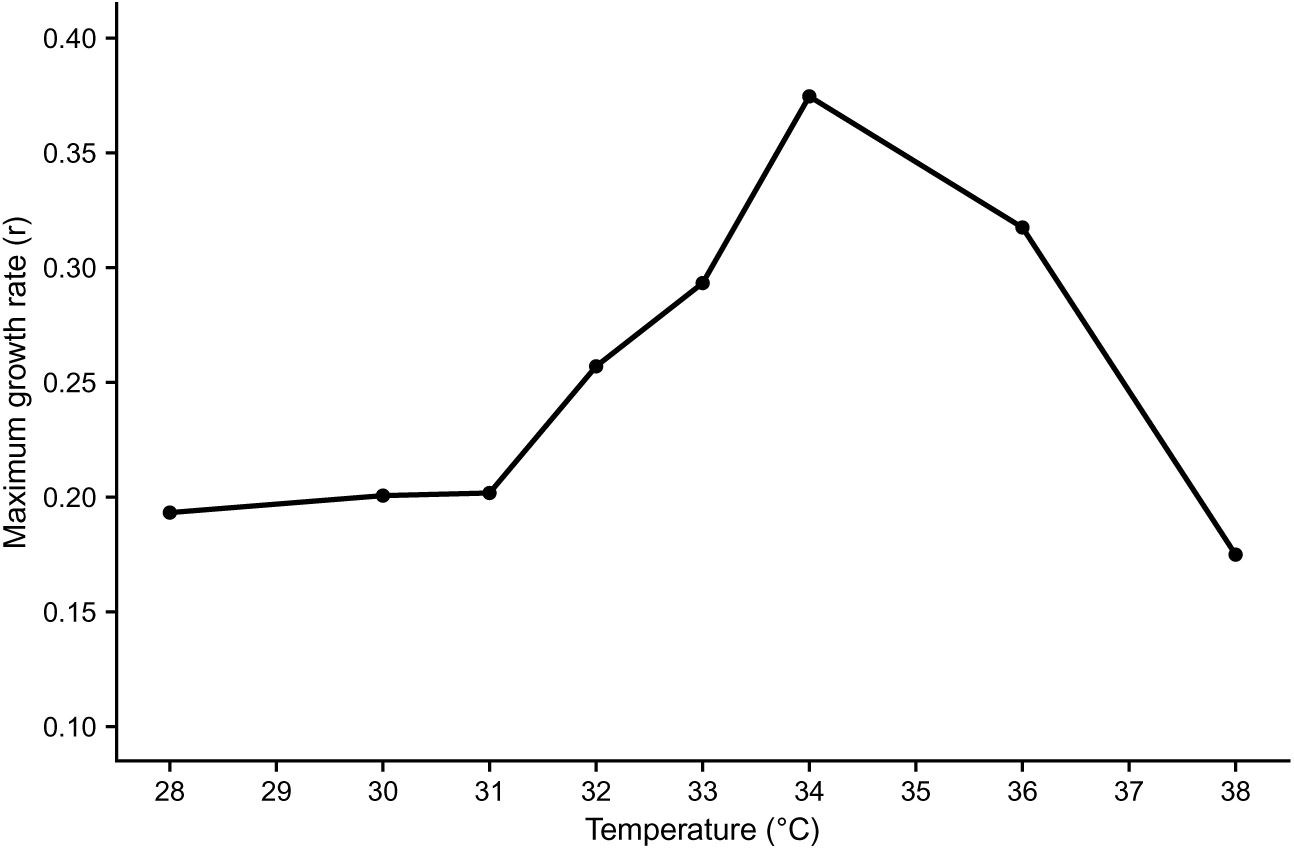
Thermal performance curve of the ancestor genotype *h*^+^ *ade6^M^*^216^.

We grew the experimental populations in Edinburgh minimal medium supplemented with adenine and uracil. The large populations were cultured on 24-well plates with a total volume of 2 ml in each well, and every 24 h 300 µl of culture was transferred to 1.7 ml of fresh medium. The small populations were grown on 96-well plates that contained 250 µl per well, and every 24 h 40 µl of culture was transferred to 210 µl of fresh medium. The transferred volumes were chosen to optimize the speed of evolution i.e. minimize the chances that rare beneficial mutations are lost in population bottlenecks (Wahl *et al*., 2002). The temperature treatments were generated in growth chambers (MTM-313 Plant Growth Chamber, HiPoint Corp., Taiwan) in which the well plates were incubated in programmable compartments. In each plate, we had control wells containing only medium to monitor cross contamination. In addition, on the 96-well plates, the wells on the edges were filled with water to avoid evaporation.

During the experiment, the number of generations was estimated by measuring the optical density of the cultures between transfers with temperature-controlled spectrophotometers (Bioscreen C®, Oy Growth Curves Ab, Ltd., Finland). The evolution experiment lasted for 6 months, and the estimated total number of generations was 464 for the small populations, and 507 for the large populations. After the experiment replicates of the evolved populations were frozen in 40 % glycerol at −80 ^◦^C to be used later in later experiments. Also parts of the samples were plated on low adenine plates to see if contamination had happened during the transfers. Based on the colour of the colonies, some populations were contaminated with cells from other samples and were excluded from the following experiments. However, in all cases the contaminated populations represented only one replicate per strain and per treatment, so there were still three replicates left of those combinations.

### 2.3 Competition experiments

To test adaptation to the evolutionary environment, and possible trade-offs between alternative environments, we measured the competitive fitness of the evolved strains. We defined the trade-offs as concurrent adaptation to one temperature treatment and resulting maladaptation to an alternative treatment. The evolved strains competed with an ancestor of the same mating type but a different *ade6* allele in their evolution environment and in the mean temperature environment. Strains evolved in mean temperature competed in all competition environments. Competitions at constant temperatures lasted five days. Competition in fast fluctuations lasted five days, including four cycles, and in intermediate fluctuations 13 days, including one cycle. Because the cycle length of slow fluctuations was 40 days, the strains evolved in this treatment were competed at constant mean and extreme temperatures. Each competition assay had three replicates, which equalled 1080 competitions in total.

Before competition, the strains were taken out of the freezer to defrost. All strains were first grown separately in liquid culture in a shaking incubator at 150 rpm at 30 ^◦^C. After 48 hours of growth, the optical density of the culture was measured to estimate the right cell concentration of the competition mix. The two competitors were mixed with an estimated ratio of 1:1 and the sample was plated on a low adenine plate to count the initial ratio of colonies at the beginning of the competition (*t*_0_), and again at the end of the experiment (*t*_1_). The bicultures were grown on 96-well plates containing 230 µl medium and 20 µl of cell suspension. The temperature treatments were created in thermal cabinets (Lab Companion, ILP-12; Jeio Tech, Seoul, Korea), and cultures were transferred to fresh medium every 48 hours.

The colony counting plates (*t*_0_ and *t*_1_) were incubated for five days at 30°C, after which they were photographed and the number of colony forming units (CFU) were counted manually for the evolved strain and the ancestor. The relative fitness of the evolved strain relative to the ancestor was calculated as the change in proportions of the colonies between platings (*t*_0_ and *t*_1_). The trade-offs were defined to occur when evolved strains had higher fitness relative to the ancestor in one temperature treatment and lower fitness in another treatment.

### 2.4 Clone measurements

Based on the results of the competition experiments we investigated the growth parameters, maximal growth rate (*r_max_*), and carrying capacity (*K*), for individual clones from the large populations and the ancestors. We studied the mechanism of adaptation, trade-offs, and how much variation is maintained in the populations. First, we created a clone libraries by plating samples from the 80 evolved large populations and isolated single colonies. Four clones were sampled from each population for a total of 320 clones. The clones were first grown for 48 h to a high density in 1.5 ml cryotubes, then 200 µl of the culture was mixed with 200 µl of 80 % glycerol on a 100-well Bioscreen plates (Bioscreen C®, Oy Growth Curves Ab, Ltd., Finland) and frozen at −80 ^◦^C. The clones of the evolved populations were randomized on four plates, and four replicates of each ancestor population were also included on each plate, as well as wells with only medium to control for contamination.

To measure growth parameters for the clones, we used the Bioscreen spectrophotometer (Bioscreen C®, Oy Growth Curves Ab, Ltd., Finland), which measures optical density at 600 nm every 5 min on 100-well plates. The frozen clone libraries were cryoreplicated to a new Bioscreen plate that contained 300 µl of fresh medium, and this plate was incubated at 34 ^◦^C for two days before starting the clone measurements. The optical density was measured to control for the initial densities in analyses, and 2 µl of culture was transferred to a new Bioscreen plate with fresh medium. Then growth of the clones was monitored for 4 days, and growth curve data obtained from each well. The measurements were made at 30 ^◦^C, 34 ^◦^C, 38 ^◦^C, and at fast temperature fluctuations, for which the Bioscreen was programmed to first set the temperature to 30 ^◦^C for 10 min, then increase the temperature to 38 ^◦^C at the rate of 0.2 °C*/*h, hold at 38 ^◦^C for 10 min, and then go back to 30 ^◦^C at the same rate. This makes one temperature cycle last for about 14 h, which is faster than in the evolution experiment, but the exact same cycle could not be replicated on Bioscreen for technical reasons. Each plate was measured four times in all of the temperatures, giving four replicates per clone per temperature.

### 2.5 Statistical analysis

#### 2.5.1 Competition experiments

First, we calculated the relative fitness for the evolved populations in each competition assay. Following Hartl and Clark (1997), the relative fitness of two asexually growing strains has the following relationship to their ratios over time

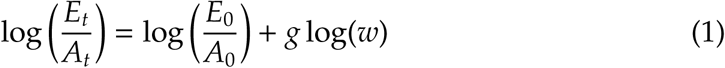

where *E_t_* is the CFU of the evolved population at time *t*, *A_t_* is the CFU of the ancestor at time *t*, *g* is the number of generations separating initial and final time points, and *w* is relative fitness of the evolved population relative to the ancestor. Solving for *w* yields the relative fitness of the evolved population in the competition assay. To examine how different treatments affected adaptation, we fitted a Bayesian generalized linear mixed model using Hamiltonian Monte Carlo algorithm, implemented in Stan (Carpenter *et al*., 2017). Stan was interfaced from R by using the brms package (Bürkner, 2017). The model was

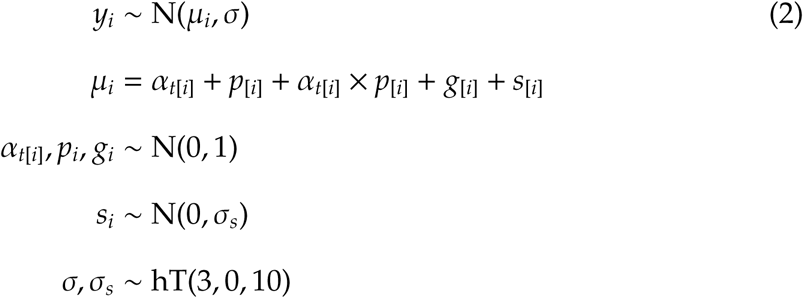

where *y_i_*was the relative fitness for *i*th competition assay, *α_t_*[*i*] was the temperature treatment, *p_i_* the population size, *g_i_* the genotype of the ancestor, and *s_i_* the identity of the evolved strain, which was included in the model as a random factor. Priors for standard deviations were the half-location scale version of Students’s t-distribution, with 3 degrees of freedom, location 0 and scale 10. This corresponds to a weakly informative prior.

The model included the effects of the temperature treatment, population size, their interactions, the effect of the ancestor, and the random effects of the replicates within a strain. The evolution environment and the competition environment were combined into a one index variable (temperature treatment) with 12 categories. The identity of the evolved strains means the specific replicate population from the evolution experiment. The ancestral genotype included combinations of the mating type and the *ade6* allele. The relative fitness data was standardized before running the model. We used four chains, with 1000 iterations of warmup and then 1000 iterations of sampling. We monitored the convergence of the MCMC chains using the *R*^^^ statistic. No convergence problems were observed. We considered the effects to be statistically different from zero when their 95 % highest posterior density intervals (HPDI) did not overlap with zero. We run also separate models for the small and large populations to be able to compare the relative fitness between different competition environments and population sizes by calculating the posterior estimates for the differences.

#### 2.5.2 Clone measurements

For the clone data, we first estimated *r_max_* and *K* from the growth curve data with the growthcurver R package (Sprouffske and Wagner, 2016). After a manual inspection of the growth curve fits, some of the clones were found to grew poorly at certain temperatures and result in unrealistically high estimates for *r_max_*and *K*. The values for these clones and were set to 0.01, and the wells that had no growth at all were marked as a missing data. For the strains that did not reach the stationary phase in four days, the value of *K* was replaced with the maximum optimal density in the data. After obtaining the growth curve parameters for each sample, we estimated environmental, genetic, and population variance components. For this the model was

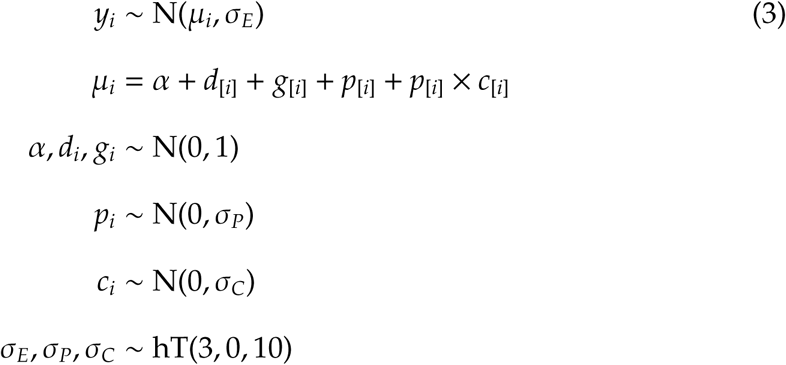

where *y_i_* was either the maximum growth rate or the carrying capacity of *i*th growth curve, *α* was the intercept, *d_i_* was the density on pregrowth plate, *g_i_* was the ancestor genotype, *p_i_* was the population identity and *c_i_*the clone identity. The environmental standard deviation was *σ_E_*, the population standard deviation was *σ_P_*, and the clone standard deviation was *σ_C_*. The variance components were obtained by squaring the standard deviations. By parameterizing the model in this way, the clone results were nested within a population. We fitted this model for all combinations of evolutionary environments and assay environments. We used four chains, with 1000 iterations of warmup and then 8000 iterations of sampling with thinning set to two. No convergence problems were observed. All statistical analyses presented were performed with RStudio (R version 4.2.2).

## 3 RESULTS

### 3.1 Competition experiments

#### 3.1.1 Adaptation to temperature

When we examined if the populations had adapted to the evolutionary treatments, we observed that clear adaptation had happened only in the populations that evolved at extreme temperature of 38 ^◦^C (Figure 2). For these populations fitness had increased 12 % relative to their ancestor and the estimate of their relative fitness was 1.12 [1.05, 1.19] (posterior mean and 95 % HPDI). No significant adaptations were observed for populations evolved at constant mean or at fluctuating temperatures (Figure 2). Yet, we did observe that the evolution in a particular temperature treatment caused fitness costs in other treatments. The populations that evolved at fast temperature fluctuations had 9 % lower fitness at the mean temperature 0.91 [0.83, 0.97], and populations that evolved at mean temperature had 28 % lower fitness at the extreme temperature compared to their ancestors 0.72 [0.65, 0.78] (Figure 2). However, no trade-offs were observed between competition environments, as the evolved populations with a significant adaptation or maladaptation did not have a reciprocal fitness increment or decrement compared to the ancestor in alternative temperature treatments (Figure 2).

**Figure 2:**
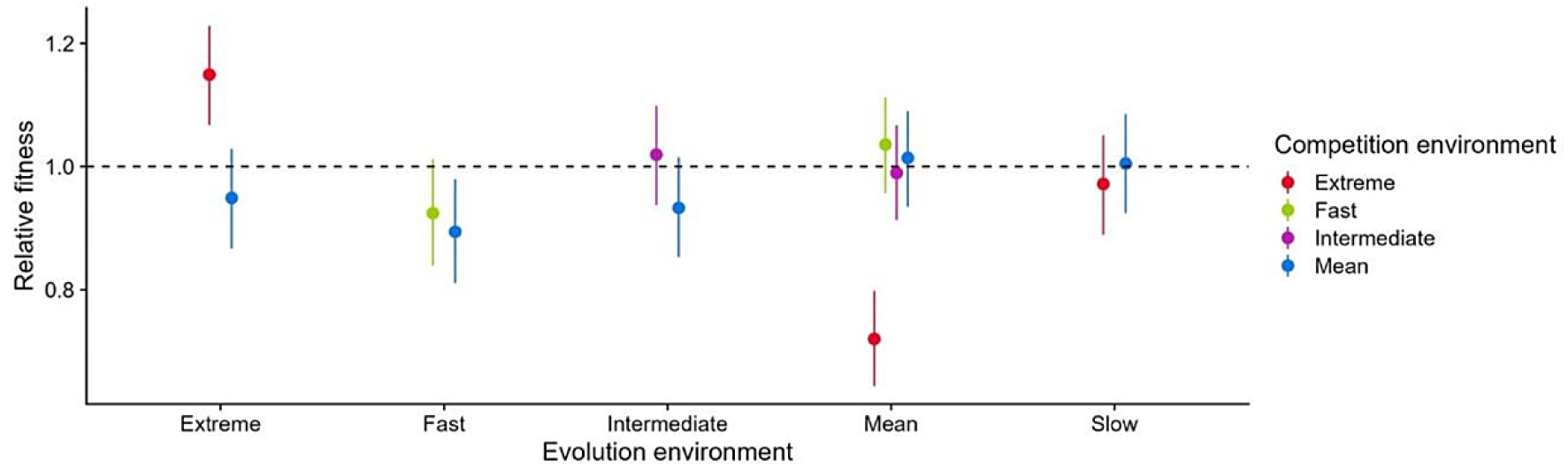
The relative fitness of the evolved strain compared to the ancestor over all populations, shown as a posterior mean with 95 % HPDI. The dashed line is the ancestor fitness set to one.

#### 3.1.2 Population size effects

Next we tested whether the larger populations adapted more efficiently to temperature treatments and whether the population size affected the occurrence of trade-offs in constant and fluctuating temperatures. The effect of population size was non-significant when tested with a model over all populations −0.03 [−0.11, 0.04]. When we run the separate models for the population sizes, there were some treatments in which the large populations seemed to do better than the smaller ones (Figure 3). Hence, we tested the differences between population sizes separately for each evolution and competition treatment (Figure 4). The difference between large and small populations was significant when populations evolved and competed at high extreme temperature 0.14 [0.01, 0.27] (Figure 4A). There was also a significant difference between large and small populations when they evolved at mean temperature and competed at fast fluctuations 0.14 [0.02, 0.27] (Figure 4B).

**Figure 3:**
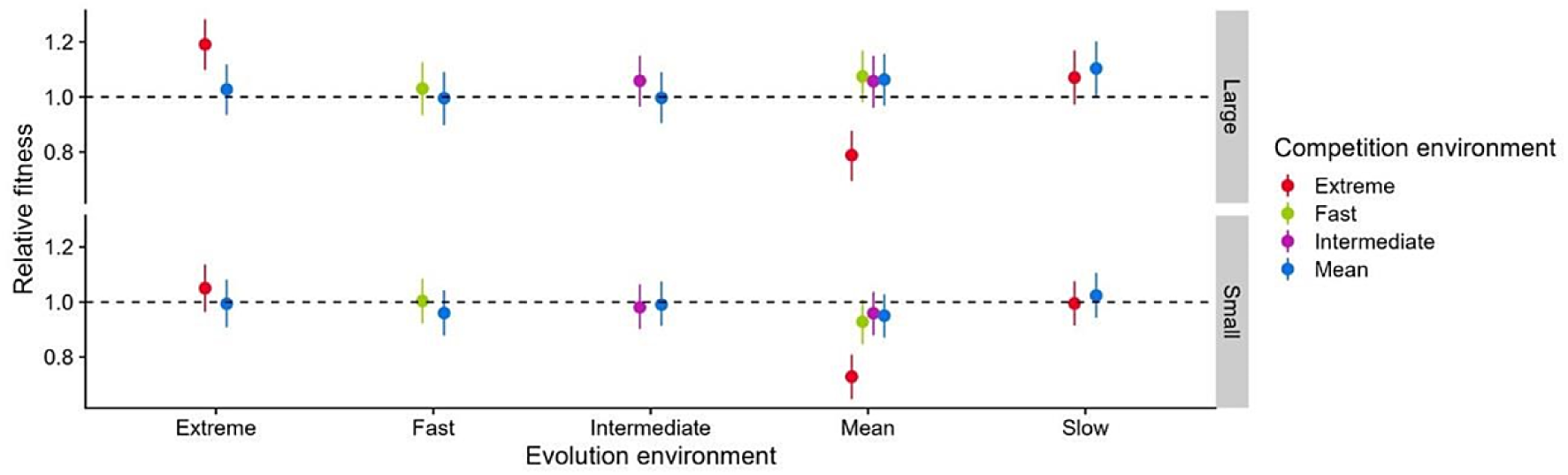
The relative fitness of the evolved populations compared to their ancestors for data divided by the population size. The relative fitness is shown as a posterior mean with 95 % HPDI. The dashed line is ancestor fitness set to one.

**Figure 4:**
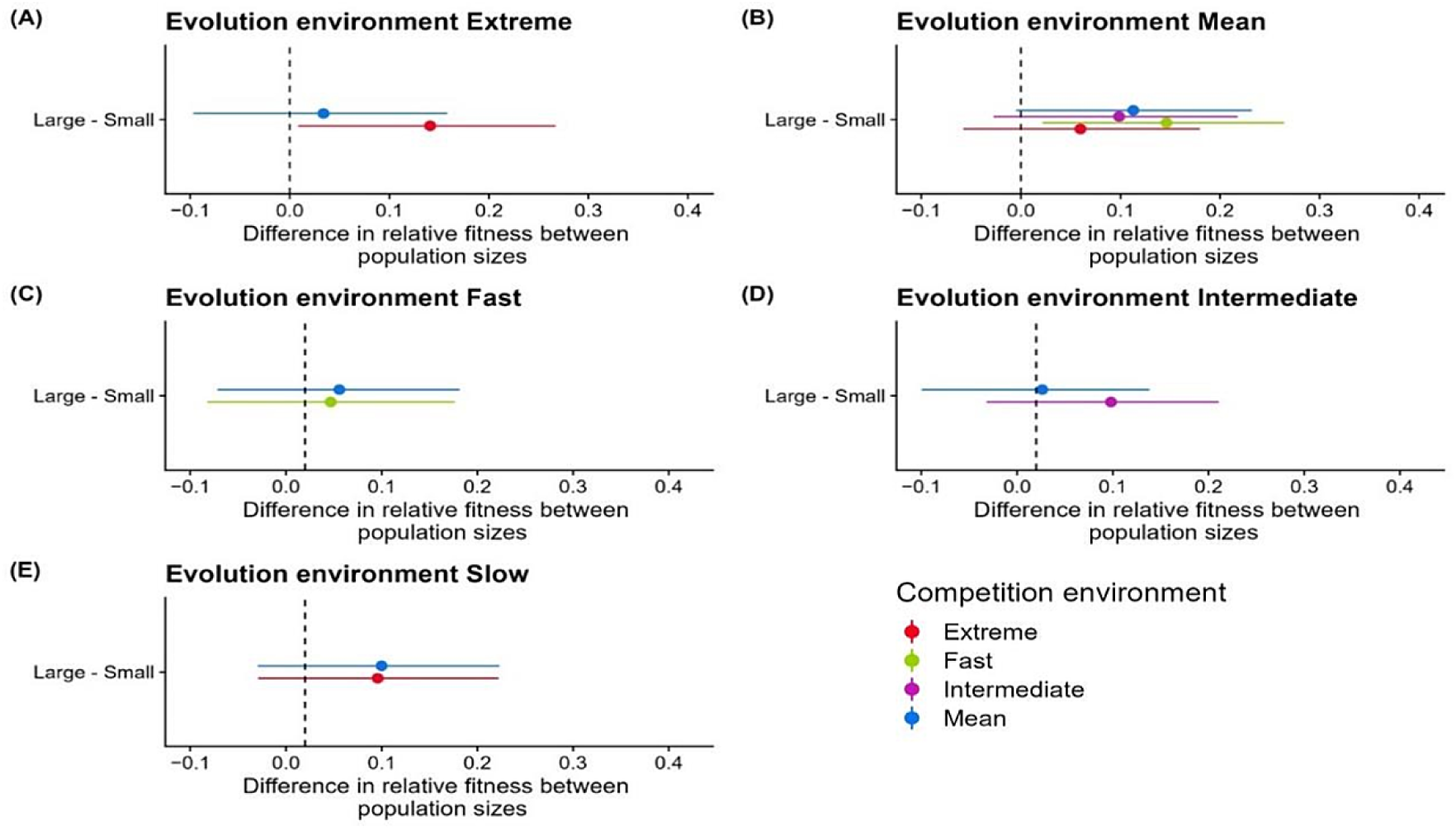
The difference in relative fitness between population sizes at different temperature treatments. The differences are statistically significant when the 95 % highest posterior density (HPDI) does not overlap with zero.

Then we examined if the relative fitness was higher at temperatures that the populations evolved compared to the mean temperature (Figure 5). We found that both large and small populations that evolved at high temperature were better competitors in a matching competition environment than in an alternative mean competition environment, with estimates of 0.16 [0.12, 0.22] and 0.06 [0.01, 0.10] respectively (Figure 5A). Similarly, large and small populations that had evolved at constant mean temperature were better competitors in a matching competition environment than in an alternative extreme competition environment, estimates 0.27 [0.22, 0.33] and 0.22 [0.18, 0.26] respectively (Figure 5B). Furthermore, we observed that the large populations that evolved at intermediate fluctuations were slightly better competitors in a matching competition environment than in an alternative mean competition environment 0.06 [0.005, 0.12] (Figure 5D). No differences in relative fitness were observed for populations that evolved at fast or slow fluctuations (Figure 5C and E).

**Figure 5:**
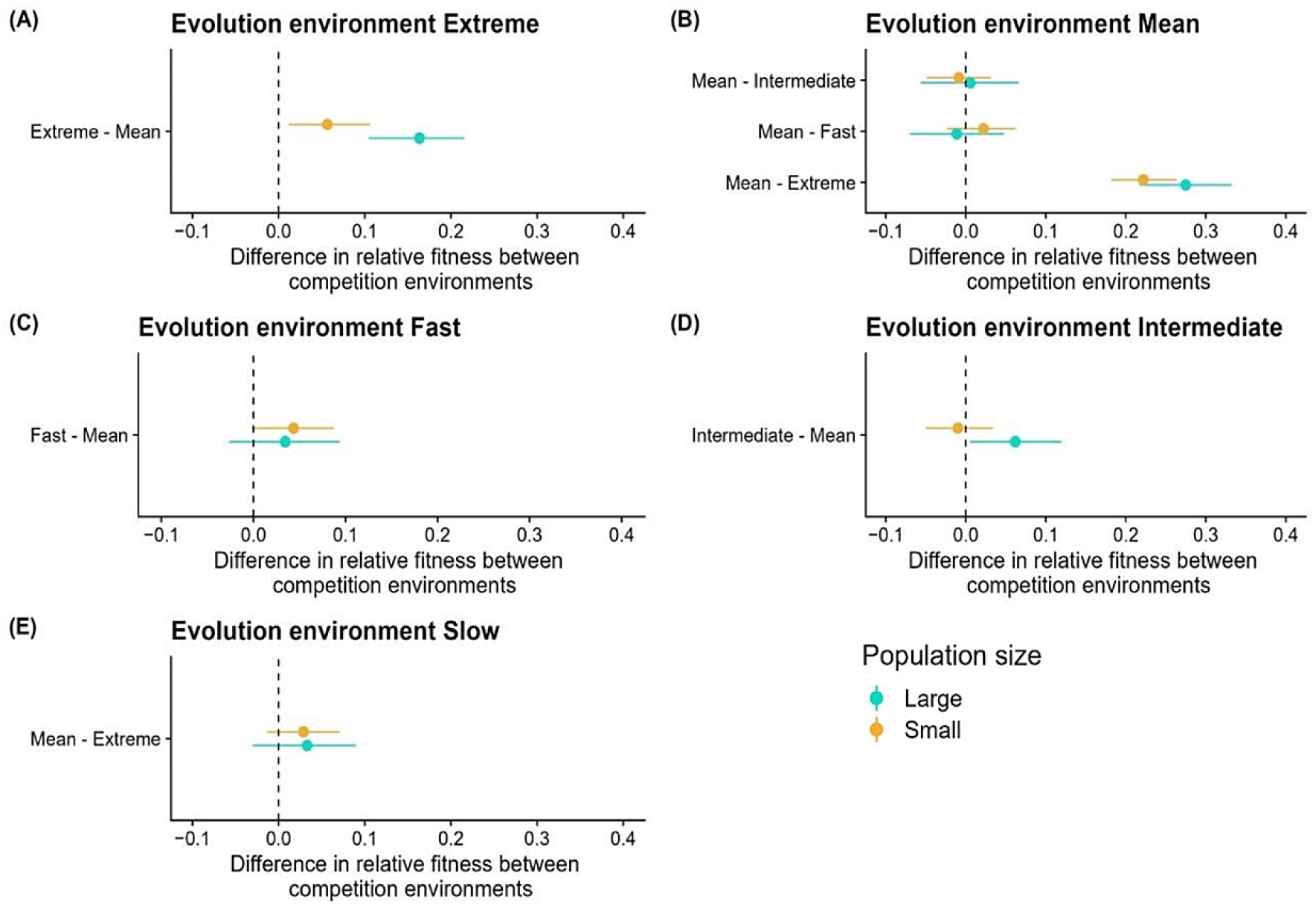
The difference in relative fitness between competition environments for large (green) and small (yellow) populations. For the statistical significance, the 95 % highest posterior density (HPDI) does not overlap with zero.

### 3.2 Clone measurements

#### 3.2.1 Changes in growth parameters

The growth rate had most consistently increased at extreme temperature for the clones that evolved at extreme temperature (Figure 6), which was in line with our observations from the competition experiments. We observed also that growth rate at 30 ^◦^C and at 34 ^◦^C had increased for the clones that evolved at mean temperature or at fluctuating temperatures (Figure 6). However, the growth rate at 38 ^◦^C had a more variable response, as it had decreased in many clones, but there were also cases where growth rate had increased (Figure 6). For the carrying capacity, the only consistent increases at 38 ^◦^C were observed for the clones that evolved at extreme temperature (Figure 7). Otherwise the carrying capacity had decreased at 38 ^◦^C for the clones that evolved at mean temperature or at fluctuations (Figure 7).

**Figure 6:**
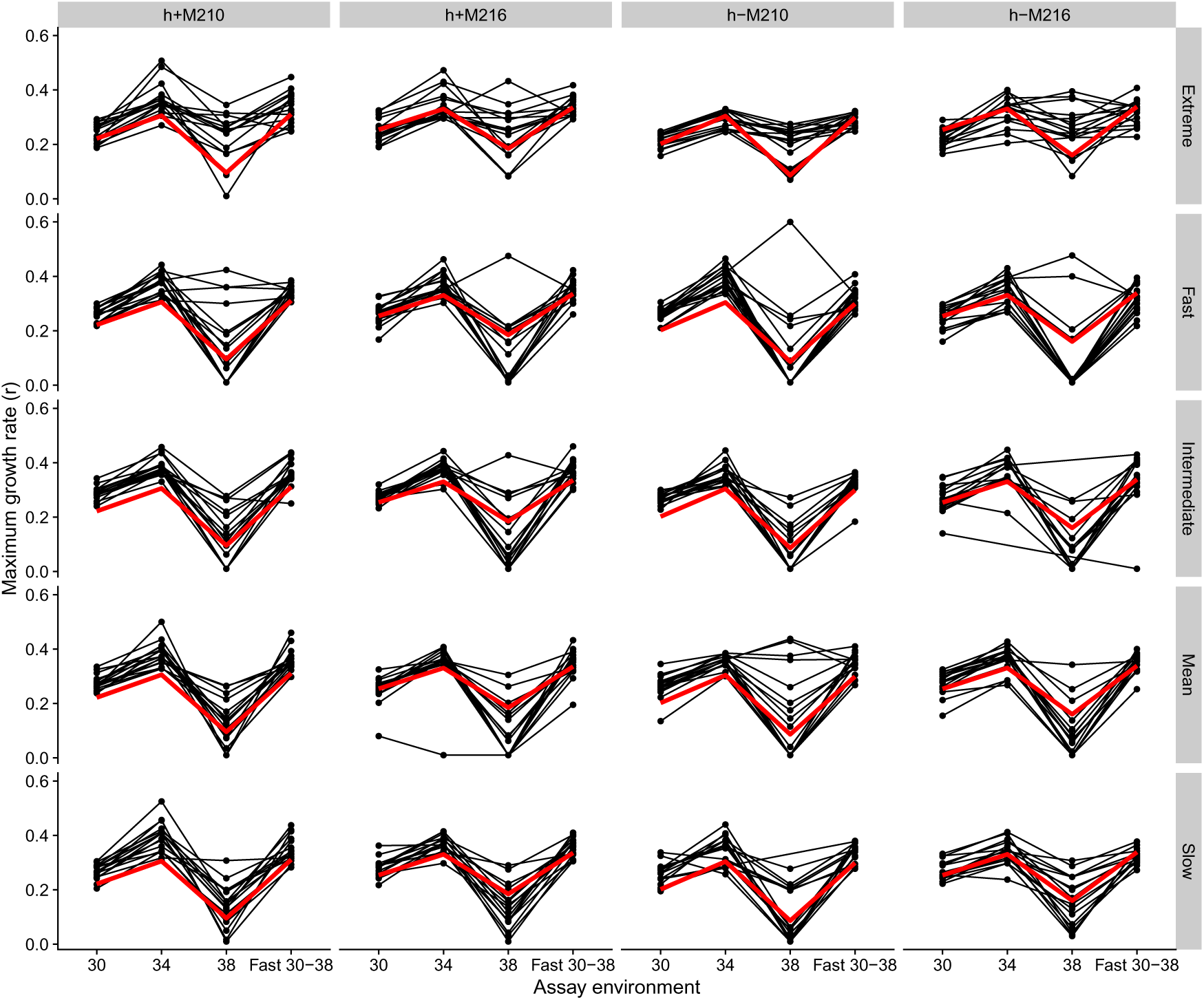
The maximum growth rates (*r_max_*) of clones sampled from the large populations. For each clone, the measurements were made at different constant temperatures and at fast fluctuations. The evolutionary treatments are on rows, and columns show the ancestor of the evolved clones.

**Figure 7:**
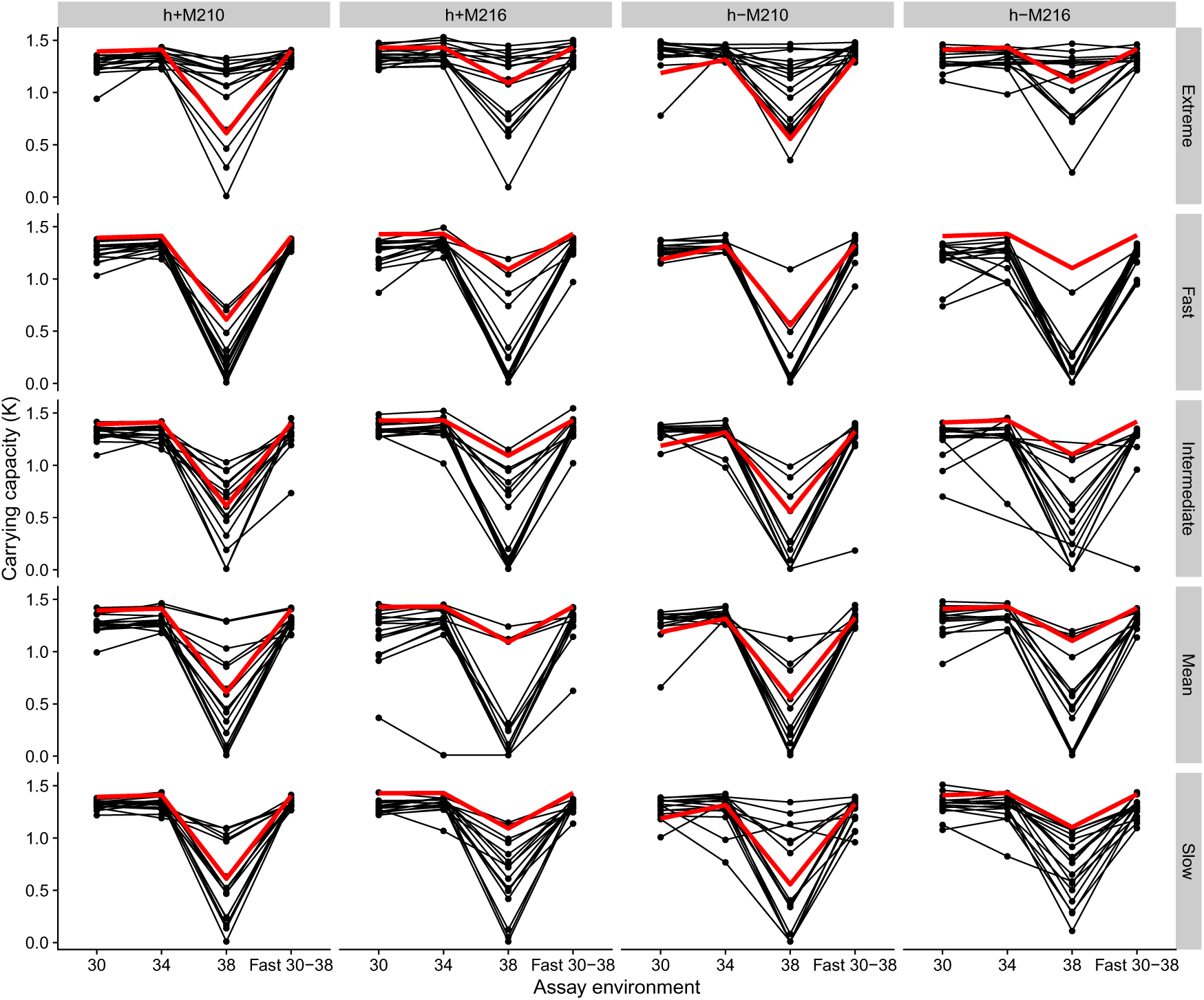
The carrying capacities (*K*) of clones sampled from the large populations. For each clone, the measurements were made at different constant temperatures and at fast fluctuations. The evolutionary treatments are on rows, and columns show the ancestor of the evolved clones.

#### 3.2.2 Genetic, environmental and population variances

To understand if fluctuating temperatures maintain more genetic variation, or if there are indications of bet-hedging or genetic canalization, we examined the genetic, environmental and population variances for the clones (Figure 8). For the genetic variance we observed that constant temperatures seemed to maintain more variation than fluctuating temperatures, as the populations evolved either at extreme or mean temperature had the largest amount of genetic variation for growth rate (Figure 8A), and carrying capacity (Figure 8B). Furthermore, the genetic variance in growth rate and carrying capacity at 38 ^◦^C was high in all populations (Figure 8), whereas the variances at 34 ^◦^C were high only in populations that evolved either in constant 38 ^◦^C or constant 34 ^◦^C. Thus, we found no support that fluctuating environments would maintain more genetic variation.

**Figure 8:**
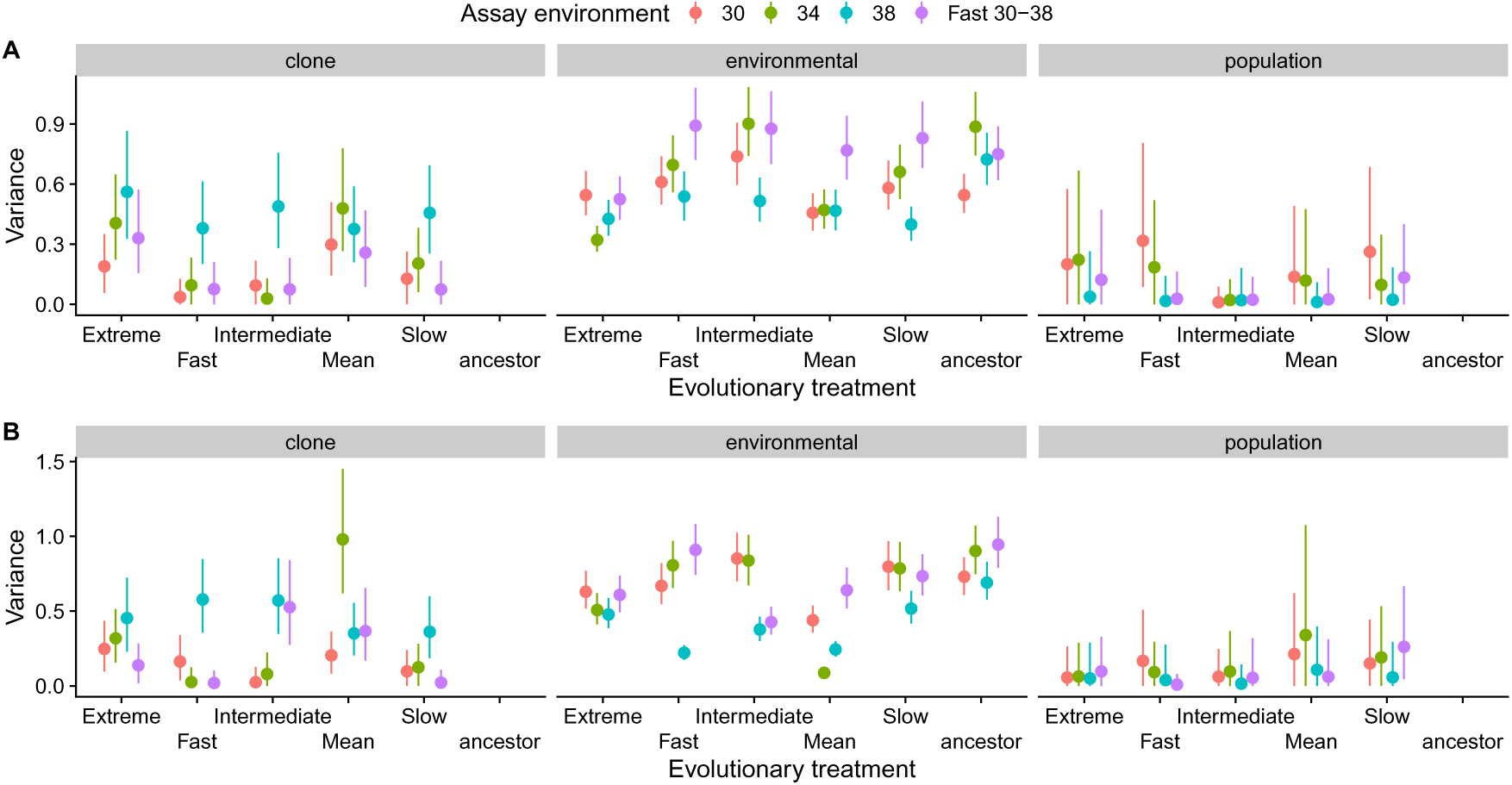
Clone, population and environmental variances for A) maximum growth rate (*r_max_*), and B) carrying capacity (*K*), at different temperature treatments. On x-axis are the evolution environments and different colours show the assay environments (°C). The variance among clones is equal to genetic variance.

The environmental variance had gone down from the ancestor level for both growth rate and carrying capacity in clones that evolved at constant 34 ^◦^C when measured at the same temperature (Figure 8). This happened also to a lesser extent for populations that had evolved at constant 38 ^◦^C. In contrast, for the populations that had evolved at fluctuations, the environmental variances were more similar to the ancestor (Figure 8). These results indicate that the evolution at constant temperatures tended to make phenotypes more canalized, whereas no such changes were observed for the populations evolved in fluctuating environments. In general, the population variances had large uncertainties (Figure 8). For growth rate the largest population variances were observed when clones were assayed at 30 ^◦^C (Figure 8A), suggesting that the genetic changes that affect growth rate at this temperature tend to be different in different populations. For carrying capacity the highest population variances were observed for the populations evolved at mean temperature and assayed at mean temperature, but with large uncertainty (Figure 8B).

## 4 DISCUSSION

### 4.1 Adaptation to temperature

We tested the hypothesis that the evolution in constant and fluctuating environments could lead to specific adaptations (Botero *et al*., 2015; Kristensen *et al*., 2020), specifically that different types of plasticity evolve at fluctuations of varying speed. However, we only observed clear adaptation to constant heat, as only populations that evolved at 38 ^◦^C increased their fitness relative to the ancestor. Populations evolved at mean temperature did not show significant fitness improvements compared to the ancestors at constant mean, fast or intermediate fluctuations. This could mean that the ancestors were already close to their thermal optimum and well-adapted to the experimental mean temperature, but also having high levels of phenotypic plasticity that makes them tolerate fluctuations (Bideault *et al*., 2023). However, populations evolved at mean temperature clearly lacked some aspects of heat tolerance, as they had 28 % decrease in relative fitness when competing at extreme temperature. The extensive adaptation and maladaptation observed in populations that had evolved at constant temperatures indicates that these conditions can act as a stronger selective pressures than temperature variation (Bennett *et al*., 1992; Kingsolver *et al*., 2009).

In general, when populations evolved in thermally variable environments, no clear adaptations were detected to fast, intermediate, or slow fluctuations. This supports the idea that the ancestor genotypes have attributes that make them readily tolerate fluctuations, for example, yeast populations might optimize their fitness by tuning the rate of switching between individual phenotypes depending on the frequency of environmental fluctuations (Acar *et al*., 2008). On the other hand, in fluctuating environments the direction and intensity of selection varies, which could make adaptation to require more generations to occur, by slowing down the fixation and purging mutations from a population (Snell-Rood *et al*., 2010; Kingsolver and Buckley, 2017). The evolution of thermal generalists has been observed in evolution experiments made with bacteria for thousands of generations (Bennett *et al*., 1992; Kassen, 2002; Ketola *et al*., 2013), but the fitness usually evolves fast during the first thousand generations, which is closer to the length of our study. In competition experiments, the only significant maladaptation to mean temperature was found for populations that had evolved under fast fluctuations, as they were 9% weaker competitors than their ancestors. This is in line with studies suggesting that tolerance to fast fluctuations requires allocation to dynamic stress response, that is costly at nonstressful conditions (Sørensen *et al*., 2003; Ketola and Saarinen, 2015). Interestingly, for populations evolved at slow fluctuations, there was no comparable maladaptation to extreme temperature as with populations that had evolved at mean temperature. Slow fluctuations had an approximated cycle of 100 generations which should be increasingly experienced as constant conditions, and hence, genetic adaptation be favored as an adaptive mechanism over plasticity and leading, for example, to evolution of two distinct specialist lines (Botero *et al*., 2015; Kronholm, 2022).

On average, we found low between population variance for both *r_max_*and *K*, suggesting that populations with the same thermal history tended to evolve similar phenotypes due to uniformity of selection pressures (Ketola and Kronholm, 2023). Overall, most of the variation for both growth parameters was attributed to environmental variation between clones. This suggests that, in general, the high thermal sensitivity of growth parameters to evolutionary treatment could stem from bet-hedging or phenotypic plasticity (Tonsor *et al*., 2013). Since clone libraries were randomized and the conditions were controlled during measurements, we can rule out that this observation would originate from a systematic error caused by external factors such as differences in handling the samples. In general, the populations from fluctuations had an equal variance with ancestors, whereas populations from constant temperatures had evolved lower environmental variance. This contradicts the idea that alternating temperatures in fluctuating environments would lead to higher environmental variance, affecting the accessibility of genetic variation to selection (Ketola and Kronholm, 2023). However, populations from constant temperatures showed genetic canalization that prevents evolutionary change by hiding genetic variation from selection due to long lasting selection pressure and thus makes phenotypes less sensitive to changes in temperature (Kawecki, 2000). It has been suggested that ancestral plasticity is more likely to lead to genetic canalization in constant environments where plasticity is costly (Scheiner and Levis, 2021).

The genetic variation within populations did not differ much between evolutionary treatments for either *r_max_* or *K*. However, there was an indication of more genetic variance within populations that had evolved at constant temperatures. This is opposite to the common expectation that fluctuating environments would maintain higher amount of genetic variation, if alleles take longer to fix due weaker selection or antagonistic pleiotropy. However, the evidence also from other experiments has been mixed (Bürger and Gimelfarb, 2002; Kassen, 2002). Interestingly, there were high levels of genetic variance in the ability to grow at extreme temperature within populations from different evolution environments. This supports the extensive evolutionary response to extreme temperature that was found in competition experiments. We observed also a high genetic variance in *K* for populations evolved at mean temperature, which is probably explaining the significant maladaptation to extreme heat in competition experiments. From reaction norms we were also able to detect that the extreme-evolved populations tolerated heat better than their ancestors, as they had higher *r_max_*and *K*. In competition experiments, populations evolved at mean temperature had drastic maladaptation to extreme heat, but the reaction norms showed that still a fraction of the clones grew better than their ancestor in extreme environment. Similar results were found in an experimental evolution experiment showing that most, but not all bacterial lines adapted to low temperature were maladapted to high temperature (Bennett and Lenski, 2007).

In our experiment, the populations originating from fast, intermediate, and slow fluctuations seemed to do on average worse than their ancestors at 38 °C. It has been shown in a meta-analysis, that if organisms evolve to be more plastic, there will be a trade-off with fixing higher upper thermal limits (Barley *et al*., 2021). However, the evolution under thermal fluctuations of different frequency improved populations growth rate at fast fluctuations and at more optimal temperatures 30 °C and 34 °C. This indicates that thermal fluctuations could select for better ability to grow fast during the times of more benign conditions, for example, by reversible plasticity (Kristensen *et al*., 2020). The same was not detected for the carrying capacity, as populations did generally worse than their ancestors in all treatments. This could mean that, due to cost associated with a fast growth rate in short-term, the population density could be lower in long-term (Bideault *et al*., 2023).

### 4.2 Population size e**ff**ects

Chavhan *et al*. (2021) have suggested that large populations that evolve at fluctuating environments could escape fitness costs in alternative local environments, whereas small populations would experience more trade-offs. On the other hand, when populations evolve at constant temperatures, larger populations should have more trade-offs in alternative environments (Chavhan *et al*., 2020). In addition, Chavhan *et al*. (2021) demonstrated that large bacterial populations can avoid fitness costs in alternative environments due to the enrichment of beneficial mutations in the same generalist line. In their literature review, Chavhan *et al*. (2021) also pointed to several studies that showed evidence for an indirect link between population size and environmental variability affecting fitness trade-offs (Bennett and Lenski, 1999; Buckling *et al*., 2007; Ketola and Saarinen, 2015).

We attempted to test the hypothesis of Chavhan *et al*. (2021), but did not observe any consistent differences in trade-offs for large and small populations. Both small and large populations that had evolved at constant mean temperature were worse competitors in the extreme competition environment than their ancestors. The only significant difference between population sizes was found when large populations had evolved at intermediate fluctuations, as they were slightly better competitors in a matching environment than in alternative mean temperature. We did observe that larger populations had higher relative fitness at extreme temperature, but no differences were observed for the other evolutionary treatments. A previous experimental evolution study with yeast has also found that populations with larger effective population size have more extensive adaptive responses to stressful conditions (Samani and Bell, 2010). It is also likely that since there was no evolutionary adaptation to the fluctuating temperatures in general, no differences were observed between the different population sizes.

Another alternative for why we did not detect a significant population size effect in general, could be that the effective population sizes chosen for the evolution experiment were too similar. The literature review of bacterial experimental evolution studies form Chavhan *et al*. (2021), suggested that in fluctuating environments, populations with *N_e_* ≈ 10^8^ are less likely to show fitness costs than smaller populations with *N_e_*≤ 10^7^. In our experiment, the difference was smaller due to technical limitations, but the same order of magnitude (small *N_e_* = 10^6^ and large *N_e_* = 10^7^). However, when Chavhan *et al*. (2021) tested the hypotheses concerning populations size and environmental variability in an evolution experiment with bacteria, the difference between population sizes they had was even larger. Another possibility regarding our experiment is that population size can have both positive and negative effects on fitness, which could cancel out each other. For example, even though higher mutational supply is beneficial in large populations, simultaneously occurring beneficial mutations can compete for fixation, slowing down evolution by clonal interference (Gerrish and Lenski, 1998). Conversely, small populations are less likely to get beneficial mutations, but the evolution could be accelerated due to faster fixation rates (Handel and Rozen, 2009).

## 5 CONCLUSIONS

In a summary, our results supported the idea that extreme heat forms a strong selection pressure, leading to prevalent adaptive responses. Also, if populations are maladapted to cope with high temperatures, there will be drastic fitness costs. Fluctuating temperatures did not seem to exert similarly high selection pressure to evolve plasticity. Along with this, other studies have suggested that high extreme temperatures can select for larger shifts in TPCs than the changing mean temperatures (Angilletta Jr *et al*., 2010; Buckley and Huey, 2016). However, we did not find strong evidence for trade-offs or expected differences in adaptation to constant and fluctuating temperatures. One plausible explanation is that ancestors were already near the fitness optimum at experimental mean temperature and at fluctuations spanning over mostly favorable temperatures. Furthermore, trade-offs may be rare just because genetic correlations between different temperatures for both constant and fluctuating conditions tend to be positive (Moghadam *et al*., 2020; Räsänen *et al*., 2024). Based on variance components, we were also able to conclude that evolution under constant temperatures maintained more genetic variation and led to lower environmental variance, indicating that genetic canalization can make phenotypes less sensitive to changes in temperature.

## Acknowledgements

This study was funded by the University of Jyväskylä Doctoral Programme in Biological and Environmental Science (ER), and grants from Emil Aaltonen foundation (ER), and the Academy of Finland (IK: 274769 592 and 321584).

## Data access

These data and scripts used for the analyses will be deposited to XXXXX under accession number YYYYY.

